# Lamin A/C directs nucleosome-scale chromatin remodeling to define early lineage segregation in mammals

**DOI:** 10.64898/2026.01.01.696913

**Authors:** Alice Sherrard, Liangwen Zhong, Caroline Hoppe, Srikar Krishna, Scott Youlten, Curtis W. Boswell, Stephen Cross, Fiona E. Sievers, Goli Ardestani, Denny Sakkas, Liyun Miao, Zachary D. Smith, Berna Sozen, Antonio J. Giraldez

## Abstract

Chromatin organization underlies gene regulation and cell fate specification, yet how nucleosome-scale chromatin structure contributes to lineage segregation during early development remains unknown. Here, we resolved chromatin ultrastructure during the first lineage decision in mouse and human, which forms the pluripotent inner cell mass (ICM) and trophectoderm (TE). To achieve this, we developed dual-tilt chromatin electron tomography (2T-ChromEMT) that allows multiscale visualization of chromatin architecture. Our analysis reveals that TE cells of both species display denser chromatin with nucleosome aggregation at the nuclear periphery. We show upregulation of the nuclear matrix protein Lamin A/C within the TE lineage across mouse, human, and opossum embryos, indicating that its regulatory role is conserved across eutherian and marsupial species. Loss of Lamin A/C reduces heterochromatin at the nuclear lamina in TE cells, reactivates pluripotency genes, and impairs mouse blastocyst expansion and human blastoid formation. These findings define the nucleosome-resolution chromatin signatures of early mammalian lineages and establish Lamin A/C–mediated chromatin organization as a conserved mechanism in the exit from pluripotency and maintenance of trophectoderm identity.

---

**T**he first lineage decision in mammalian development establishes the inner cell mass (ICM) and trophectoderm (TE). These two unique cell lineages display restricted developmental potentials—the ICM exhibits pluripotency and forms the future fetus, and the TE generates the cells that will develop into the placenta^1–3^. Spatially defined epigenetic and transcriptional programs underlie these cell identities. They are initiated *de novo* after fertilization and are reinforced by mechanically regulated Hippo signaling^4–7^, where the nuclear matrix protein Lamin A/C has been implicated in the coupling of mechanical forces to ICM and TE fate^8^. Pluripotency is established with expression of *Nanog, Sox2*, and *Pou5f1 (Oct4)* in ICM precursor cells situated towards the embryo’s interior. In outer cells that form the TE, *Cdx2* and *Gata3* repress the pluripotency factors^1,3,9–12^. However, whether Lamin A/C directly regulates chromatin structure to drive these gene expression profiles remains unknown.

While epigenetic marks associated with these cell states have been described^13^, how chromatin conformation promotes and stabilizes ICM and TE identities remains unclear. This raises the fundamental question of whether distinct chromatin structural properties define the first lineages in mammalian development. Species-specific differences in chromatin organization between mouse and human pluripotent stem cells^14–18^ highlight the diversity of chromatin states underlying pluripotency. In addition, Hippo-independent lineage specification in marsupials^19^ points to alternative regulatory mechanisms driving and maintaining early cell fate decisions. Together, these findings underscore the need for comparative analyses to understand how chromatin architecture drives lineage specification. Such questions have been difficult to address due to limited access to human embryos at different developmental stages. However, recent advances in stem cell-based human blastoids, which mimic blastocyst-stage organization, provide a tractable model to investigate these regulatory events *in vitro*^20–22^. Another challenge lies in the technical limitations of existing approaches for analyzing chromatin structure, which spans orders of magnitude. No current imaging or genomics method captures all scales with high fidelity from a single sample.

Although subnucleosome-scale visualization of chromatin structure has been enabled by chromatin electron tomography (ChromEMT)^23^, new acquisition and quantitative analysis approaches are needed to connect nucleosome organization with higher-order chromatin structure for multiscale studies in mammalian embryo models.

Here, we adapted ChromEMT^23^ to uncover how chromatin is remodeled during the earliest lineage segregation in mouse and human. We identified cell type-specific chromatin signatures that are conserved across species, as well as defining characteristics of human embryonic chromatin. We linked specific features of TE chromatin to the conserved upregulation of Lamin A/C, which we show is essential for blastocyst formation in mouse and human models. Our findings define Lamin A/C as a conserved driver of the first cell fate decision across mouse, human, and opossum embryos.

## RESULTS

### 2T-ChromEMT resolves multiscale chromatin ultrastructure *in situ*

To define how chromatin structure changes during the first cell fate decision in mammalian embryos, we required an integrated imaging approach that allows the analysis of *in situ* chromatin structure across scales. We based our approach on ChromEMT^23,24^, which uses photo-oxidation of the DNA dye, DRAQ5^23^, to produce chromatin-specific EM contrast (see methods, Fig. 1a, S1a). However, ChromEMT previously lacked accessible imaging strategies and methods to analyze chromatin. To reconstruct 3D chromatin structure across scales in early embryos, we thus developed dual-tilt (2T) electron tomography followed by isotropic reconstruction using a neural network model^25^. This approach, which we call 2T-ChromEMT, corrects the missing wedge artifact observed in tomography, while using widely accessible electron microscopy equipment and machine learning chromatin segmentation. This enables global analysis of individual chromatin chains and nucleosomes throughout the nuclear volume, selecting nucleosome densities of 11 nm +/−3 nm for analysis (the size of a nucleosome^26^, Fig. 1b, S1b, S1d, Movie S1-2). For our integrated approach, we developed quantitative imaging tools that quantify various structural features across several parameters, including chromatin density, nucleosome position, nucleosome distance, and chromatin diameter. Chromatin density was measured by quantifying chromatin volume within a 40 nm sphere across the tomogram to generate contour maps (Fig. S1c), while nucleosome packing was measured by quantifying the distance to 10 nearest neighbors. Together, 2T-ChromEMT enables multiscale characterization of chromatin remodeling during differentiation, from chromosome compartments to chromatin loops and individual chains at sub-nucleosome resolution (1.61 nm / pixel, Fig. 1b).

**Figure 1.**
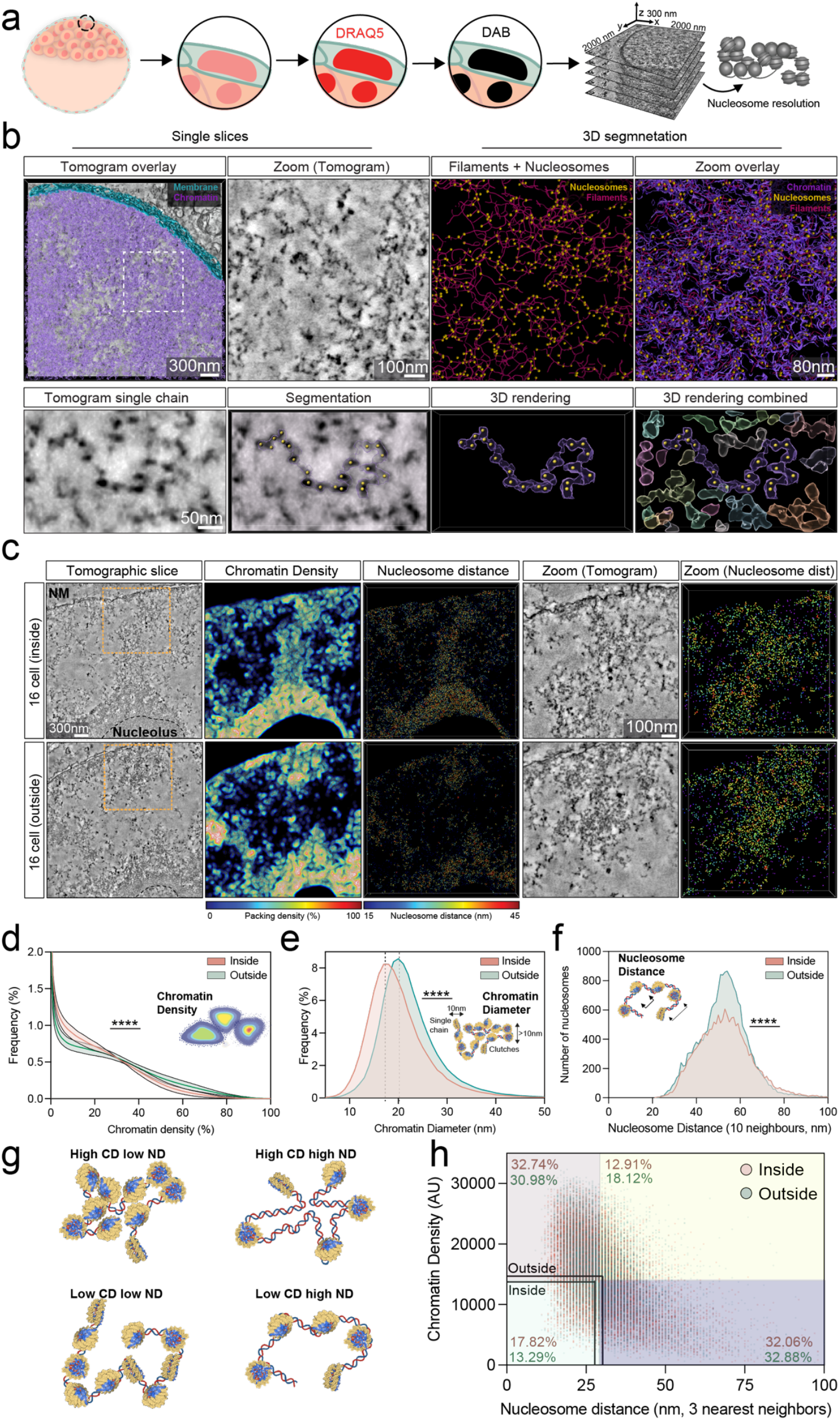
Quantification of chromatin structure across genomic and functional scales. a, Cartoon of 2T-ChromEMT protocol. Mouse blastocysts are stained with DRAQ5 and photo-oxidized in the presence of diaminobenzidine (DAB). DAB polymers deposit onto the chromatin and are bound by osmium tetroxide to provide DNA-specific EM contrast. Embryos are cut into 300 nm sections and 2µm x 2µm nuclear regions are imaged by dual-axis electron tomography to achieve sub-nucleosome resolution. b, Segmentation of a mouse 16-cell embryo. First panel shows an overlay of chromatin segmentation in purple and a single tomographic slice. Nuclear membrane in cyan. Next panels are zooms of the region indicated by the white box, segmentation mask is shown in purple, chromatin traces in pink, and nucleosomes in yellow. Below shows segmentation and 3D rendering of a single chromatin chain (purple) with nucleosomes (orange). Right shows neighboring chains in different colors. c, Tomograms and quantitative maps of 16 cell embryo inside and outside cells, zoomed region is indicated by the orange box. Chromatin density, contour maps displaying chromatin packing density indicated by the color-coded scale bar. Nucleosome distance, nucleosomes colored based on their distance to 3 nearest neighbors in 3D as a proxy for nucleosome clustering, indicated by the colored scale bar. d, Quantification of chromatin packing density, error bars show standard deviation. e, Quantification of chromatin diameter. f, Quantification of the distance of nucleosomes to 10 nearest neighbors. d-f, N= 5 inside cells, 8 outside cells, 2 biological replicates. p****<0.0001, Wilcoxon test. g, Cartoon representing different organizations of chromatin density (CD) and nucleosome distance (ND). h, Quantification of CD and ND in inside and outside cells of a 16-cell mouse embryo. Colored boxes indicate above and below the median chromatin and nucleosome density. Numbers indicate the percentage of chromatin in each of the four quadrants for each cell type.

As an initial comparison of chromatin architecture in cells of distinct developmental states, we first examined undifferentiated mouse embryonic stem cells (mESC) and differentiated mouse embryonic fibroblasts (MEF). We found that mESCs were distinguished by a decondensed chromatin structure that is organized within large regions of interchromatin space, with further decompaction observed at the scale of individual nucleosomes. At the global scale, MEF cells showed uniform chromatin density (median of 31.4%), consistent with previous results^23^. In contrast, mESCs showed a significant decrease in chromatin density (median 21.2%) with 23.4 % of the nucleus largely devoid of chromatin (<10% density, Fig. S1e-f), presumably occupied by nuclear bodies or RNA molecules^27,28^. These results were consistent across multiple biological replicates (n=6 cells, Fig. S1g). At the nucleosome scale, we observed a decrease in internucleosome distance in differentiated MEF cells, with nucleosomes positioned on average 6 nm closer than in mESCs (Fig. S1h). These results are consistent with the known formation of larger, more dense nucleosome clutches and higher levels of chromatin compaction upon differentiation^29–34^. Surprisingly, chromatin fibers in mESCs exhibit significantly increased diameters, with 14.4% measuring above 30 nm (Fig. S1i). Such emergence of large chromatin diameters resembles higher-order folding models previously proposed for 30 nm fibers^35–37^. These fibers are absent in differentiated cells^23,38^, but are present in pluripotent stem cells, where globally decondensed chromatin may require higher-order organization. Together, we show that 2T-ChromEMT resolves chromatin architecture at subnucleosome resolution and enables a quantitative framework to distinguish structural organization across scales in cells at different stages of differentiation

### Global chromatin reorganization primes the trophectoderm fate in 16-cell mouse embryos

To determine when chromatin structure diverges during early lineage specification, we examined mouse embryos at the 16-cell stage, when the pluripotent inner cell mass (ICM) and trophectoderm (TE) lineages are defined^3,6^. They can be distinguished by their positions within the embryo, with the ICM precursors situated internally and the lineage-restricted TE externally^5–7^. We found that both ICM and TE precursor cells were characterized by large regions of the nucleus exhibiting low density (~35.8% of the nucleus had <10% density; Fig. 1c-d) and pronounced perinucleolar chromatin compaction (Fig. S1j). This distribution was markedly different from that observed in MEFs and mESC, while displaying a more similar distribution to mESCs (Fig. S1e-f). Yet, outer lineage-restricted TE cells could be distinguished by an increase in high chromatin density (Fig. 1d), thicker chromatin fibers (median diameter 22.1 nm vs. 20.2 nm for inner cells; Fig. 1e), and denser nucleosome clusters (Fig. 1f). Together, this suggests early structural priming of the TE lineage, consistent with higher chromatin compaction upon differentiation.

Because direct visualization of chromatin structure *in situ* enables multiscale chromatin analysis, we also determined the relationship between chromatin density and nucleosome distance, which are generally thought to be anticorrelated (Fig. 1g). Genomic methods such as Hi-C^39^ and ATAC-seq^40^ measure these properties separately, and they have shown that transcriptionally active regions are accessible at the nucleosome level and occupy low-density euchromatic compartments. Consistent with this, we found that 32.5% of the chromatin in ICM and TE cells of 16-cell embryos had low chromatin density (below the median <14758 AU) along with high nucleosome distance (above the median >28 nm), while 31.9% of the chromatin had high chromatin density combined with low nucleosome distance, representing accessible and compact chromatin states, respectively. However, a subset of regions deviated from these canonical states. We observed a population of multiple chromatin chains that were close together with wide nucleosome spacing (compacted yet accessible chromatin; 12.91% and 18.12% in inner and outer cells, respectively), reminiscent of spatial co-activation of genomic loci ^41,42^. Conversely, we also observed a population with fewer chromatin chains but tightly packed nucleosomes (de-compacted inaccessible; 17.82% and 13.29% in inner and outer cells, respectively) (Fig. 1g-h). These observations suggest that chromatin accessibility and compaction are not strictly coupled in early mouse embryos, revealing subnuclear heterogeneity in chromatin architecture (Fig. 1h). This is reminiscent of the heterogeneity we detected in chromatin density distribution within TE precursors, which exhibited a significant decrease in low density chromatin and a concomitant increase in high density chromatin compared to the ICM (Fig. 1d). Together, these results indicate that the first lineage specification begins with spatially regulated chromatin compaction, with fibers hyper-condensed at the nuclear periphery and nucleolus, interspersed with expansive interchromatin space. The TE lineage is thus structurally primed for differentiation, showing early compaction at both chromatin levels.

### Peripheral chromatin compaction is a conserved hallmark of TE differentiation

To determine whether lineage-specific chromatin features are conserved across species, we compared the ICM and TE of mouse blastocysts and human blastoids (Fig. 2a, S2a). In mice, ICM and TE displayed similar chromatin densities and diameters (median density, 25.5% and 22.7%; diameter, 19.3 and 19.4 nm; Fig. S2b-c). However, TE cells displayed increased subnuclear heterogeneity in chromatin packing, including higher levels of condensed and chromatin-free regions compared to the ICM, indicating the formation of dense chromatin compartments (Fig. S2b). This distribution was in stark contrast to ICM and TE precursor cells in 16-cell embryos, with cells of the blastocyst exhibiting minimal nucleolar compaction and significantly less chromatin-free area. By contrast, human TE cells exhibited greater global compaction (29.4% versus. 25.5% in ICM), thicker chromatin fibers (22.2 versus 19.5 nm), and the appearance of chromatin fibers of >30 nm (Fig. S2d-e), consistent with higher-order chromatin folding^23,38^. At the nucleosome scale, both human ICM and TE cells showed a significant difference in the organization compared to that of mouse cells, with small dense nucleosome clusters sparsely distributed throughout the nucleus (Fig. 2a-b), suggesting more accessible chromatin. This may underlie the increased developmental plasticity of human blastoids compared to mouse embryos of a similar stage^43,44^. Together, these data reveal that lineage-specific chromatin architectures emerge across mouse and human embryos, with TE cells exhibiting higher compaction in both species.

**Figure 2.**
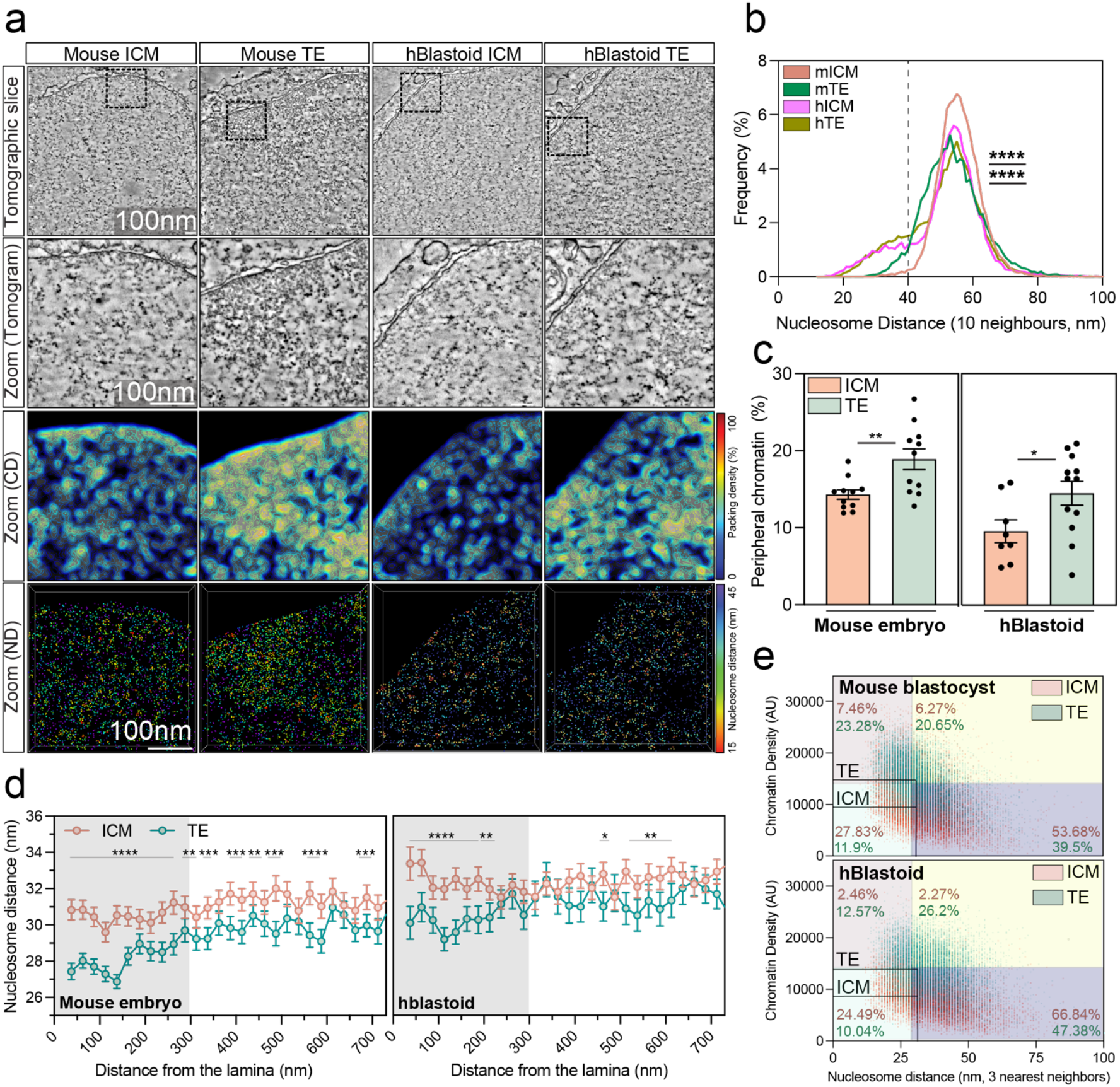
TE cells exhibit increased chromatin density and nucleosome clustering at the nuclear periphery. a, Representative images and quantitative maps of inner cell mass (ICM) and trophectoderm (TE) cells from mouse blastocysts and human blastoids imaged by 2T-ChromEMT. Top panels, single tomographic slice of the indicated cell type, zoom below of the region indicated by the black box. Third row panels, topographic density maps where chromatin density (CD) is represented as a color-coded density map. Bottom panels, nucleosome segmentation and distance (ND) coded by color. Nucleosome color indicates proximity to the 3 nearest nucleosomes in 3D. b, Quantification of the distance of nucleosomes to 10 nearest neighbors. N=7 mouse ICM and TE cells, N=9 human blastoid ICM and TE cells across four embryos and two biological replicates. Line represents the mean value. ****p < 0.0001, mouse vs human ICM, mouse vs human TE, Wilcoxon test. Line represents the mean distance. c, Quantification of the percentage of chromatin within 1 µm distance from the nuclear periphery based on transmission electron microscopy (TEM) images. N = 11 ICM and TE cells across 4 mouse embryos from two biological replicates. N = 8 ICM cells and 12 TE cells across three human blastoids in two biological replicates. **p < 0.01, *p <0.05, Student’s t-test. Error bars show SEM. d, Quantification of the average distance to the 3 nearest nucleosome neighbors in 3D as a function of distance from the nuclear membrane. Data points were divided into 25 nm positional bins starting at the inner nuclear membrane. Data were pooled from 7 ICM and 6 TE cells from mice and 8 ICM and 4 TE cells from human blastoids from four embryos across two biological replicates. *p < 0.05, **p < 0.01, ***p < 0.001, ****p < 0.0001, multiple unpaired t-test. Data show mean ± 95% confidence interval. Gray area highlights values within 300nm of the nuclear membrane. e, Representative quantification of nucleosome distance and chromatin density with medians in the indicated cell types. Colored quadrants show the median values in a 16-cell mouse embryo projected onto the indicated cell types. Numbers show the percentage of values in each of these four quadrants per cell type.

We next analyzed chromatin structure in the context of nuclear position. Both mouse and human TE cells showed a significant increase in chromatin compaction at the nuclear periphery compared to ICM cells (Fig. 2a, 2c), which was confirmed by transmission electron microscopy (TEM, Fig. S2f). These condensed chromatin domains are reminiscent of lamina-associated domains (LADs), which are condensed chromatin regions tethered to the nuclear lamina that are associated with gene silencing^45–47^. While LADs in the ICM and TE are reportedly similar; whether nucleosomes are spatially compact at the nuclear lamina in the TE remains unknown. To address this, we quantified nucleosome spacing as a function of distance from the nuclear periphery. We found that nucleosome distances within 300 nm of the nuclear membrane were ~4 nm shorter in TE compared to ICM cells in both mouse blastocysts and human blastoids (Fig. 2d). Thus, TE cells in both species display higher levels of condensed chromatin compared to the ICM, characterized by high nucleosome and chromatin density. Combined analysis of the chromatin density and the nucleosome distance revealed that the ICM across mouse and human blastoids shows an increase in low-density chromatin with closed nucleosomes (de-compacted inaccessible chromatin) compared to TE cells (Fig. 2e). These findings suggest that the de-compacted inaccessible chromatin state is associated with pluripotency, and that non-canonical chromatin states are regulated during cell differentiation. Together, we identified peripheral chromatin compaction as a conserved hallmark of TE differentiation, which points to a potential role for nuclear architecture in directing this process.

### Lamin A/C is a conserved marker of the trophectoderm

The formation of dense chromatin at the nuclear periphery, specifically in TE cells, suggests the involvement of nuclear lamina components, which regulate signaling, compact chromatin, and repress genes at the nuclear periphery^8,45,46,48^. We found that Lamin A/C was indeed strongly upregulated in the TE of mouse blastocysts at E3.5 compared to the ICM, while Lamin B1 was uniformly expressed across both lineages (Fig. 3a-b, Fig. S3a). Quantification across stages (8-cell to E4.5 blastocysts) revealed that Lamin A/C levels rose at the 16-cell stage, when TE transcriptional specification begins, and continued to increase through late-blastocyst formation at E4.5 as observed at the RNA and protein level (Fig. 3b-c, Fig. S3b). To assess whether this pattern is conserved, we examined human embryos donated for research (thawed and analyzed at day 5/6, see Methods) and observed enrichment of Lamin A/C in human TE cells compared to ICM (Fig. 3d-e). Marsupial blastocysts (opossum, Monodelphis domestica) exhibited a similar TE-specific enrichment of Lamin A/C expression at both the protein and RNA levels (Fig. 3f-h). Together, these findings identify Lamin A/C upregulation as a conserved feature of TE differentiation across eutherian and marsupial mammals. Lamin A/C– dependent organization of peripheral chromatin may thus be an ancestral mechanism to secure extraembryonic lineage identity.

**Figure 3.**
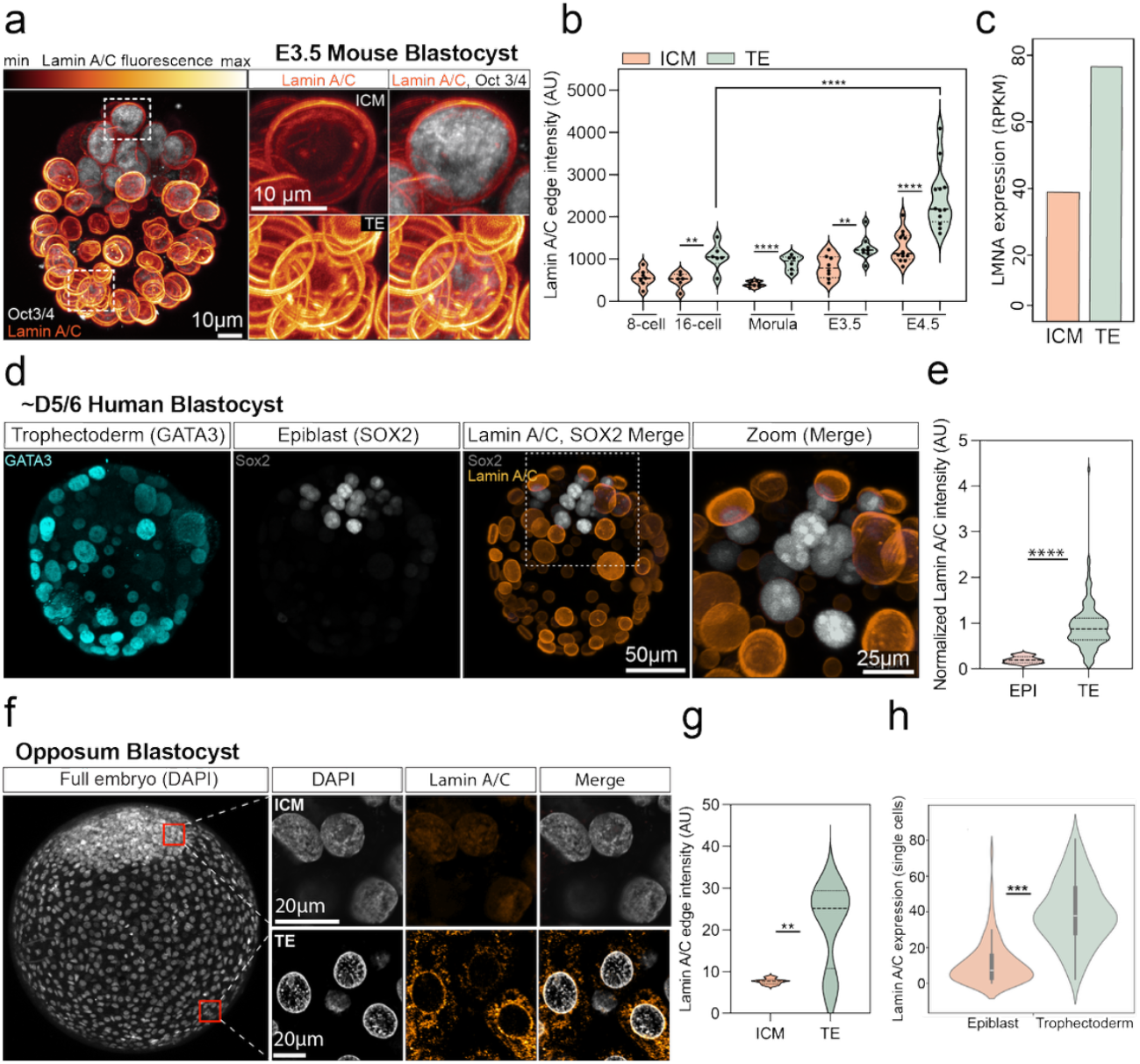
Lamin A/C is a conserved marker of the trophectoderm. a, IF images of a mouse blastocyst with LAMIN A/C (orange), OCT3/4 (ICM marker; gray). Right panels, insets outlined by the white boxes. b, Quantification of Lamin A/C nuclear edge intensity in ICM or TE cells at the indicated stages of mouse pre-implantation development. Each dot represents the median intensity of one embryo pooled across 8-273 cells per embryo; N=7 embryos (8-cell), N=6 embryos (16-cell and morula), N=8 embryos (E3.5), and N=12 embryos (E4.5), in 2 independent experiments. One-way ANOVA, **p < 0.01, ****p < 0.0001. Violin plot shows median and interquartile range. c, RNA expression (RPKM) of *Lmna* in ICM and TE cells (mean used from^64^, two biological replicates). d, IF images of human day 5/6 blastocysts obtained from Boston IVF clinic. Cyan= GATA3 (TE), gray= SOX2 (epiblast), orange= Lamin A/C. Zoom (right) highlights low levels of Lamin A/C in the ICM compared to the TE. e, Quantification of Lamin A/C intensity in cells shown in d (epiblast, EPI; and TE) normalized to DAPI. N=1 embryo; 26 epiblast cells and 128 TE cells. Student’s t-test, ****p < 0.0001. f, IF images of an opossum (Monodelphis domestica) blastocyst. Left panel shows DAPI in gray; the right panels are higher-magnificat of ICM (top) and TE (bottom) cells labeled with DAPI (gray) and Lamin A/C (orange). g, Quantification of LAMIN A/C intensity in cells shown in f (ICM and TE). N=1 embryo, >15 cells. Student’s t-test, **p < 0.01. h, RNA expression of *Lmna* from single cell sequencing of the Epi and TE of the opossum (data from^65^). ***p < 0.001, two-sided Mann–Whitney U test (Wilcoxon rank sum). p-values adjusted for multiple tests using the Benjamini–Hochberg false discovery rate.

### Lamin A/C regulates chromatin architecture and gene expression to maintain TE identity

To investigate whether Lamin A/C drives the TE-specific structural reorganization of chromatin, we generated *Lmna* KO mouse embryos using CRISPR-Cas9 (Fig. 4a). Loss of Lamin A/C protein was confirmed by immunofluorescence (Fig. 4b-c). ChromEMT revealed the abrogation of condensed peripheral chromatin in the TE, with fewer and smaller peripheral chromatin densities in *Lmna* KO TE cells (Fig. 4d-e). We further observed increased internucleosome spacing at the nuclear lamina, resulting in a configuration that resembles the chromatin organization of the wild-type ICM (Fig. 4f). In addition to a localized reduction in packing density at the nuclear lamina, depletion of Lamin A/C led to changes in chromatin organization broadly throughout the nucleus. Global packing density increased in *Lmna* KO TE versus WT cells (median density 32.9% and 22.7% respectively, Fig. S4a) with a concomitant reduction in low density chromatin in *Lmna* KO TE cells (9% vs 27.7% of the nucleus had <10% chromatin density in *Lmna* KO vs wildtype TE cells). These findings imply that in the absence of Lamin A/C, chromatin packing in the TE is more uniform, losing both hypercondensed regions at the periphery as well as chromatin-depleted regions. Accordingly, we observed a reduction in the median diameter of TE chromatin fibers (16 vs. 18.4 nm in WT) while maintaining similar internucleosome distances (Fig. S4b-c). Therefore, Lamin A/C regulates nucleosome-scale chromatin compaction at the nuclear lamina and global organization throughout the nucleus in the TE.

**Figure 4.**
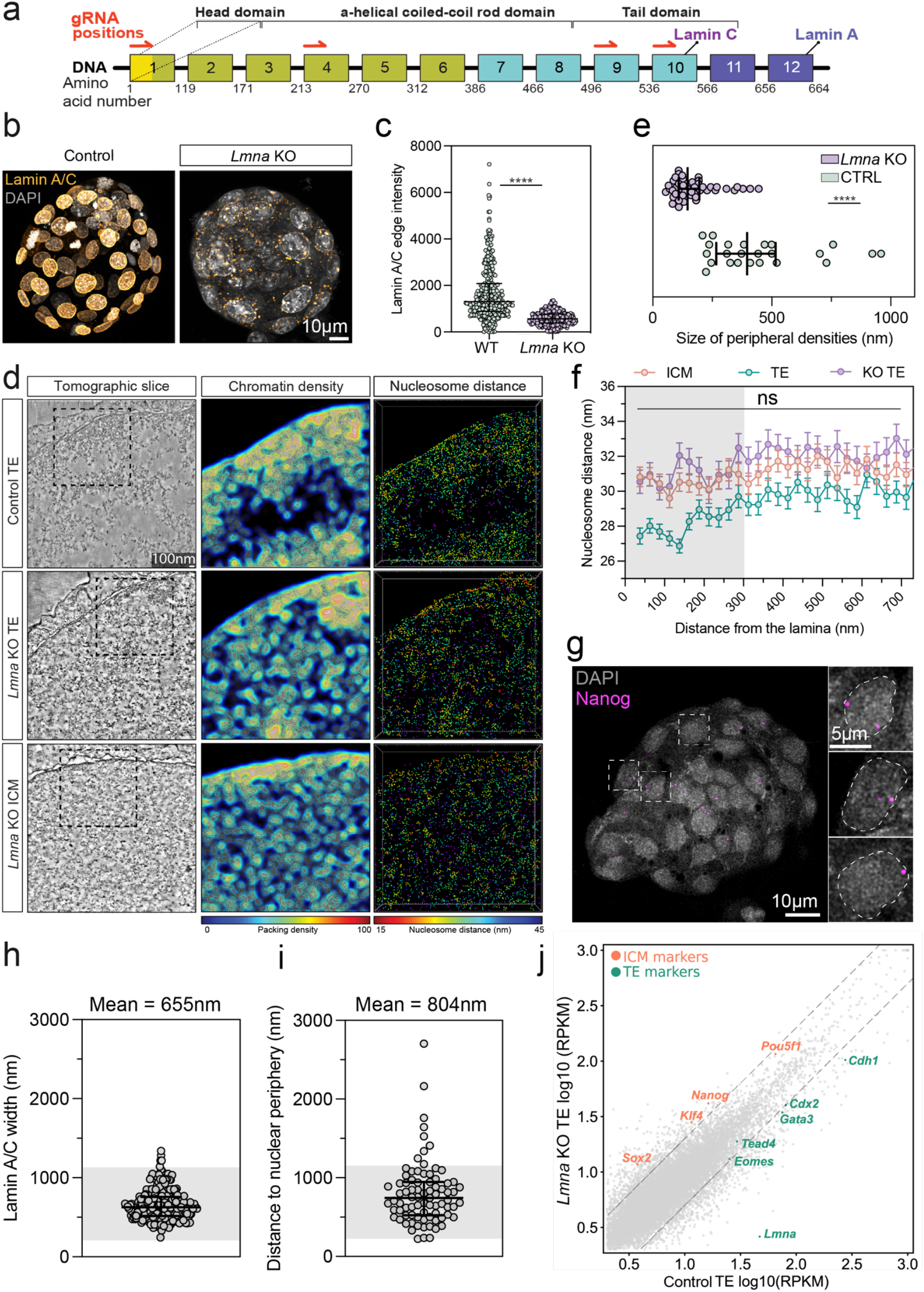
Lamin A/C regulates chromatin structure and TE gene expression. a, Schematic of the *Lmna* gene locus and the encoded protein domains. The Lamin C isoform is encoded by exons 1-9 and a small portion of exon 10, while Lamin A adds exons 11 and 12, including the CAAX box motif. Based on^66^. b, IF images of WT and *Lmna* KO blastocysts labeled with Lamin A/C antibody (orange) and DAPI (gray). c, Quantification of Lamin A/C levels in WT and KO blastocysts shown in b. N=>350 cells in 6 embryos pooled from two biological replicates. ****p < 0.0001, Mann Whitney test. Bars indicate the median and interquartile range. d, Representative tomogram (2T-ChromEMT) of a control TE cell (top), a *Lmna* KO TE cell (middle), and a *Lmna* KO ICM cell (bottom). Center and right panels are zooms (indicated by the black boxes) of the corresponding chromatin packing density maps and nucleosome distance (3 nearest neighbors) colored based on the indicated scales. e, Quantification of the length of dense chromatin regions at the nuclear periphery of the cells shown in d (control and *Lmna* KO TE cells). n=11 nuclei from 4 embryos. ****p < 0.0001, Mann Whitney test. f, Quantification of the average distance to 3 nearest nucleosome neighbors as a function of distance to the nuclear membrane. Data were binned into 25 nm sections. WT data from Fig 2d. *Lmna* KO data were pooled from five cells and statistically compared to WT ICM data. ns= non-significant, multiple unpaired t-test, data shows mean ± 95% confidence interval. g, IF image showing DNA FISH of *Nanog* in a mouse blastocyst. The right panels are zooms of the regions indicated by the dashed white boxes outlining nuclei on the left. h, Quantification of lamina thickness in Lamin A/C IF images. N=64 nuclei across six embryos in two biological replicates. Gray region highlights values in within 1.2μm of from the nuclear membrane. i, Quantification of the distance of *Nanog* loci to the nuclear periphery. N= 6 embryos, two biological replicates. Gray region highlights values in within 1.2μm of from the nuclear membrane. j, Graph showing RNA-sequencing counts from WT and *Lmna* KO TE cells. ICM marker genes are shown in orange, TE marker genes are shown in green. Dashed lines show 2-fold limits. Data is from >10 embryos in one sequencing replicate.

We next asked whether Lamin A/C-mediated chromatin organization is essential to regulate gene expression of the TE cells. Three lines of evidence support a role for Lamin A/C in repressing pluripotency genes in TE cells. First, DNA FISH revealed that both alleles for the pluripotency gene *Nanog* were consistently localized at the nuclear periphery, where they overlapped with Lamin A/C in the TE of mouse blastocysts (mean distance 804nm, Fig. 4g-i). TE-specific upregulation of Lamin A/C may therefore promote direct interactions with ICM-defining loci, leading to their repression via recruitment of known Lamin A/C interactors^49,50^. Second, we isolated ICM and TE cells from control and *Lmna* KO mouse blastocysts using immunosurgery and analyzed their lineage-specific gene expression (Fig. S4d-f). This revealed upregulation of the pluripotency genes *Nanog, Sox2*, and *Pou5f1* (*Oct4*), and downregulation of the TE markers *Cdx2* and *Gata3* in outer cells upon loss of Lamin A/C (Fig. 4j, Fig. S4e-f). Third, hybridization chain reaction (HCR) showed that these transcriptional changes occur at the single-cell level, showing increased *Nanog* and reduced *Tead4* and *Cdx2* transcripts in individual TE cells of *Lmna* KO embryos (Fig. S4g). Together, we conclude that Lamin A/C organizes perinuclear chromatin and coordinates transcriptional programs to maintain TE identity, directly linking nuclear genome organization to lineage specification in early development.

### Lamin A/C is required for TE maturation and blastocyst formation across mammals

To determine whether Lamin A/C functionally regulates TE development, we analyzed *Lmna* KO mouse embryos during preimplantation stages. Prior to TE specification (48 hours post fertilization, hpf), the TE markers YAP1 and CDX2 were expressed at similar levels between WT and *Lmna* KO embryos (Fig. S5a). However, by 72-96 hpf, *Lmna* KO embryos showed significant downregulation of CDX2 and YAP1 in TE cells (Fig. 5a-c), consistent with disrupted transcriptional programs (Fig. 4j). The loss of TE identity coincided with reduced developmental progression in *Lmna* KO embryos, with only 62% reaching the morula stage and 24% forming blastocysts, compared to 89.5% and 70% in control embryos (Fig. 5d-e). This was not due to a developmental delay, as developmental arrest persisted through 120 hpf (Fig. S5b-c). Notably, KO of *Lmna* in a single blastomere of the 2-cell embryo restored blastocyst formation (Fig. S5d), indicating that Lamin A/C acts cell-autonomously, and the WT cell can compensate for *Lmna* loss by contributing to the TE lineage. Following transfer of E3.5 blastocysts to pseudopregnant females, implantation occurred normally, but only 32.4% of *Lmna* KO embryos developed to E7.5 (vs 63.7% control; Fig. 5f, Fig. S5e). Together, these findings demonstrate that Lamin A/C regulates TE maintenance and blastocyst development.

**Figure 5.**
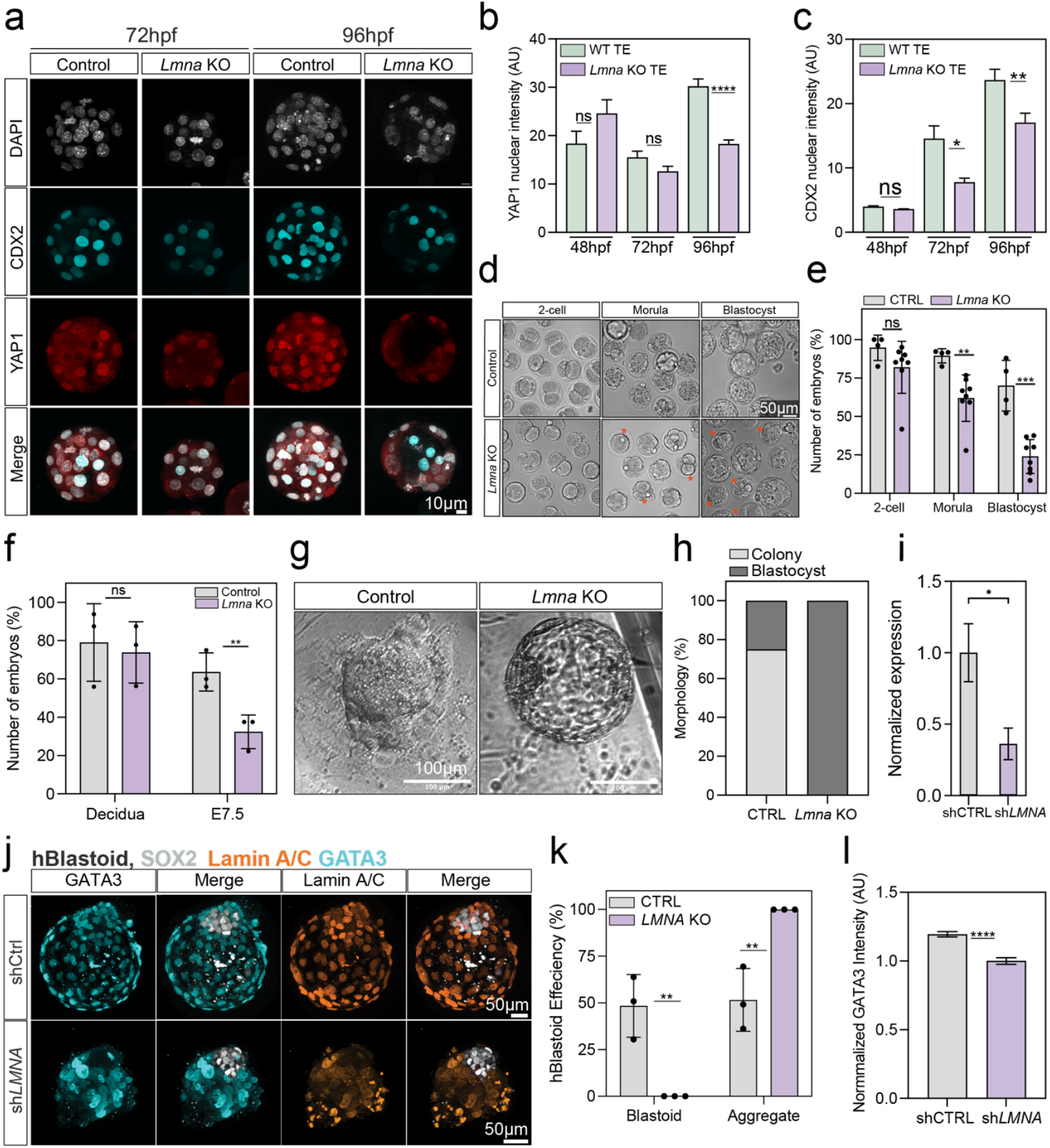
Lamin A/C regulates TE development. a, Immunofluorescence (IF) images of control and Lmna KO mouse embryos 72- and 96-hours post fertilization (hpf). CDX2 (cyan), YAP1 (red), DAPI (gray). b, Quantification of YAP1 nuclear intensity of the cells shown in a. N=8 embryos per condition, two biological replicates, ****p < 0.0001, ns= non-significant, Student’s t-test. c, Quantification of CDX2 nuclear intensity of the cells shown in a. N=6 embryos per condition, two biological replicates, Student’s t-test, *p < 0.05, **p < 0.01, ns= non-significant. d, Representative images of control and Lmna KO embryos that developed to the indicated stages. Asterisks indicate embryos with developmental arrest. e, Bar graph showing the percentage of control and Lmna KO embryos reaching the indicated stages of development. N=33 CTRL embryos and N=44 Lmna KO embryos, four independent experiments. **p < 0.01, ***p < 0.001, ns= non-significant, Student’s t-test. f, Graph showing the percentage of WT and Lmna KO embryos that formed a decidua and E7.5 embryos post-implantation. A total of 45 embryos were transferred per condition across three experiments. Each dot represents the mean of one experimental replicate. **p <0.01, ns= non-significant, Student’s t-test g, Representative images of colony progression two days post trophoblast stem cell (TSC) induction. h, Percentage of blastocysts that had formed colonies after two days post-induction (CTRL, N=8; Lmna KO, N=7). i, Bar graph showing the relative expression of LMNA by qPCR in naïve human embryonic stem cells (hESCs) treated with lentivirus expressing control non-targeting (shCTRL) and LMNA (shLMNA) shRNAs. *p < 0.05, Student’s t-test. Error bars show the standard deviation of three technical replicates in one biological replicate. j, IF images of control and LMNA knockdown (KD) human blastoids. GATA3 (cyan), SOX2 (gray), Lamin A/C (orange). Images show maximum intensity projections. k, Quantification of the number of control and LMNA KD cells that formed expanded blastoids. N=3biological replicates, N=464 CTRL blastoids, N=302 LMNA KD blastoids. Error bars show SD. **p < 0.01, Student’s t-test, 3 biological replicates l, Quantification of GATA3 intensity in shCTRL and shLMNA KD blastoids. N=10 shCTRL blastoids (1004 TE cells) and N=31 shLMNA blastoids (441 TE cells), ****p < 0.0001, Student’s t-test, 2 biological replicates. Error bars show SEM.

To further investigate the role of Lamin A/C in driving TE developmental potential, we derived mouse trophoblast stem cells (mTSCs) from WT and *Lmna* KO blastocysts. Genotyping revealed a frame shift in exon one and large deletion between exons 4-10 (Fig. S5f) that resulted in complete loss of Lamin A/C protein expression, as validated by Western blotting and immunofluorescence (Fig. S5g-h). *Lmna* KO TE cells were significantly delayed in forming TSC colonies, requiring 7-11 days compared to 2 days for wild-type TE cells (Fig. 5g-h, Fig. S5i-k), whereas pluripotent ESC derivation was unaffected (Fig. S5l). Lamin A/C thus regulates TE self-renewal and identity.

Finally, using the human blastoid model^20,21^, we tested whether Lamin A/C also regulates formation of the human blastocyst. Naïve hPSCs depleted of *LMNA* by shRNA (Fig. 5i, Fig. S5m) generated aggregates containing SOX2^+^ epiblast-like and GATA3^+^ TE-like cells but failed to form organized blastoid structures (Fig. 5j-k). LMNA-depleted aggregates also exhibited reduced GATA3 expression (Fig. 5l), reminiscent of the TE defects observed in *Lmna* KO mouse embryos (Fig. 4j). In contrast, control cells robustly formed blastoid structures with an enrichment of Lamin A/C in the TE (Fig. S5n-o). Together, these results indicate that Lamin A/C is important for blastocyst formation in human embryos, a critical stage in development that is frequently compromised in infertility

## DISCUSSION

Our study reveals that chromatin architecture is spatially reorganized to drive the first cell fate decision in mammals. We developed 2T-ChromEMT to visualize chromatin architecture at subnucleosomal resolution *in situ* during early embryogenesis and found that pluripotent ICM cells have a highly accessible chromatin structure, characterized by low chromatin compaction and dispersed nucleosomes. In contrast, lineage-restricted TE cells exhibited a defined chromatin organization at the nuclear lamina, marked by high chromatin density and closely packed nucleosomes. This structural organization is conserved between mouse embryos and human blastoids, where TE chromatin shows enhanced compaction at the nuclear periphery. We identified Lamin A/C as a conserved regulator that organizes repressed chromatin in the TE, in part by silencing pluripotency genes positioned near the nuclear lamina, such as *Nanog*. Finally, our results illustrate how 2T-ChromEMT enables *in situ* quantification of the structural and spatial features of chromatin that underlie gene regulation and reveal a conserved chromatin-mediated mechanism underlying the first cell fate differentiation.

The development of 2T-ChromEMT, combined with open-source tomography software, provides subnucleosome-resolution imaging of chromatin architecture using conventional electron microscopy. This approach complements existing genome imaging tools^51–55^ by enabling simultaneous quantification of chromatin conformation, packing density, and nucleosome spacing. By employing 2-tilt tomography, as opposed to the 8-tilt previously used in ChromEMT^23^, 2T-ChromEMT achieves isotropic reconstruction with EM equipment that is broadly accessible. With this method, we identified distinct chromatin states *in situ* characterized by variable density and internucleosome spacing. While active chromatin is classically associated with low density (“A compartments”)^39^ and wide nucleosome spacing (“ATAC peaks”)^40^, our *in situ* measurements reveal non-canonical configurations, where some loosely packed chromatin contains tightly arranged nucleosomes, and conversely, some dense chromatin exhibits wide nucleosome spacing. These states may reflect alternative modes of gene regulation, such as transcription factories, actively transcribed enhancer-promoter contacts, or regions undergoing chromatin remodeling or replication^41,42^. The relative abundance of these configurations differed between the ICM and TE, consistent with lineage-specific regulatory programs. In particular, the low density and short nucleosome distance (de-compacted inaccessible) was associated with the pluripotent ICM. This state might reflect regions that may be activated during later stages of cell differentiation or are yet to be opened by pioneer factors. Conversely, high density regions with long nucleosome distances (compacted but accessible regions) are enriched in the TE and may reflect transcriptional hubs or regions that are associated with TE differentiation. Future studies integrating transcriptional activity, histone modifications, and chromatin-associated proteins will help define the molecular functions and the regulatory potential of these distinct chromatin organizations.

By directly quantifying chromatin density at the nuclear periphery, we identified TE-specific chromatin signatures at the nuclear lamina. These dense peripheral chromatin structures may resemble lamina-associated domains (LADs) previously mapped by genomic methods in pre-implantation mouse embryos^45^. LADs have been reported to be comparable in genomic size between the ICM and TE^46^. While these genomic approaches quantify the genomic regions that are interacting with the lamina, 2T-ChromEMT directly quantifies chromatin organization at the nuclear periphery with nucleosome resolution. Our results imply that genomically defined LADs may represent a subset of a more expansive chromatin region that is in proximity to the nuclear lamina. Our results support a role for these chromatin regions in driving TE identity and blastocyst development. We further demonstrate a role for Lamin A/C in the nucleosome-scale packing of chromatin at the lamina in the TE, consistent with a recent cryo-EM study in *Lmna* KO MEF cells^47^. In addition, our data extend previous super-resolution imaging results that showed an increase of nucleosome clusters in size and density upon *in vitro* differentiation of mESCs^29^. Notably, nucleosome organization was more accessible in human embryos, characterized by smaller, denser, and more dispersed nucleosome clutches, potentially reflecting the greater plasticity of human TE cells compared to that of the mouse embryo^43^.

We find that Lamin A/C organizes repressed chromatin during the first cell fate decision specifically in TE precursors, suggesting a conserved strategy for Lamin A/C– mediated exit from pluripotency. This mechanism mirrors Lamin A/C-dependent repression of Nanog observed during *in vitro* neural differentiation of mESCs^56,57^. Beyond local repression, we found that chromatin compaction at the nuclear lamina has downstream implications throughout the nucleus. Loss of Lamin A/C resulted in global chromatin remodeling, with a reduction of hypercondensed and chromatin-depleted regions, indicating that Lamin A/C maintains global chromatin organization across the nucleus. This is consistent with genomic studies that have shown global effects in chromatin architecture upon loss of Lamin A/C during *in vitro* cardiac differentiation^57^. We thus establish Lamin A/C as a global architectural regulator that drives chromatin compaction, represses pluripotency genes and enforces lineage restriction during trophectoderm differentiation.

Lamin A/C is enriched in the TE of two eutherian mammals and a marsupial, indicating a conserved role in extraembryonic lineage specification and, by inference, placental development. Because blastocyst implantation involves large-scale biomechanical remodeling^58^, and Lamin A/C is a critical regulator of nuclear and cellular stiffness^48,59^, our findings suggest a potential link between nuclear remodeling and the establishment of the fetal-maternal interface. Lamin A/C upregulation and the repression of Hippo signaling (8-16 cell stage^6,7^) have been associated with early TE cell fate specification. However, we can derive TE cells in the absence of Lamin A/C that express TE marker genes (Fig. 5a, 5g), albeit at lower levels. We propose a two-step model where Lamin A/C regulates cellular tension to repress Hippo signaling, and also plays a direct role in chromatin compaction in TE cells that silences pluripotency genes. This is supported by our observations that pluripotency genes localize in close proximity to the nuclear lamina and that Lamin A/C is upregulated in the marsupial TE, which is independent of Hippo signaling^19^. We show that in mouse blastocysts, Lamin A/C acts as a determinant of TE fate, regulating nucleosome-scale chromatin organization and promoting lineage-specific chromatin compaction. Lamin A/C is critical for the expansion of the mammalian blastocyst, and while the phenotype is only partially penetrant, escapers might be rescued by partial redundancy with other lamina proteins such as Emerin^60^. Together, our results identify Lamin A/C as a conserved regulator that couples chromatin organization and lineage specification, promoting exit from totipotency, TE differentiation and blastocyst formation, with implications for understanding the causes of human infertility ^61–63^.

## ACKNOWLEDGEMENTS

We thank the Yale Center for Cellular and Molecular Imaging (CCMI) Confocal Facility for the use of their Zeiss 880 and Leica SP8 microscopes, as well as the CCMI electron microscopy facility for the use of the Biotwin and TF20 microscopes. In particular, we thank Kimberley Gibson and Dr. Xinran Liu for their training and support. We thank the Yale Center for Genome Engineering, particularly Dr. Tim Nottoli and Xiaojun Xing for implanting blastocyst embryos and providing facilities and expertise to train and perform this work. We thank Tova Finkelstein for assisting with E7.5 mouse embryo dissections, and Dr. Daniel Stadtmauer and Professor Gunter Wagner for providing opossum blastocysts. We further thank Micheal Murphey (Zeiss, Arivis) for helping to establish our image analysis pipelines. Finally, we thank our scientific writers Drs. Andy Cox and Linnea Weiss for their critical feedback. We further extend appreciation to Dr. Cox for his thoughtful comments and ideas throughout this project.

## AUTHOR CONTRIBUTIONS

Conceptualization and project administration, A.J.G. and A.S. Funding acquisition, A.J.G., B.S., A.S. Supervision and Resources, A.J.G., B.S, Z.S. Investigation, A.S., L.Z., L.M., C.W.B. Formal Analysis and Software, A.S., L.Z., C.H., S.G.K., S.Y., F.E.S., S.C. Visualization and Validation, A.S., L.Z., C.H. Writing, A.J.G. and A.S., review and editing from all authors.

