## Supplemental_Figures for "Lamin A/C directs nucleosome-scale chromatin remodeling to define early lineage segregation in mammals"

### SUPPLEMENTARY FIGURES

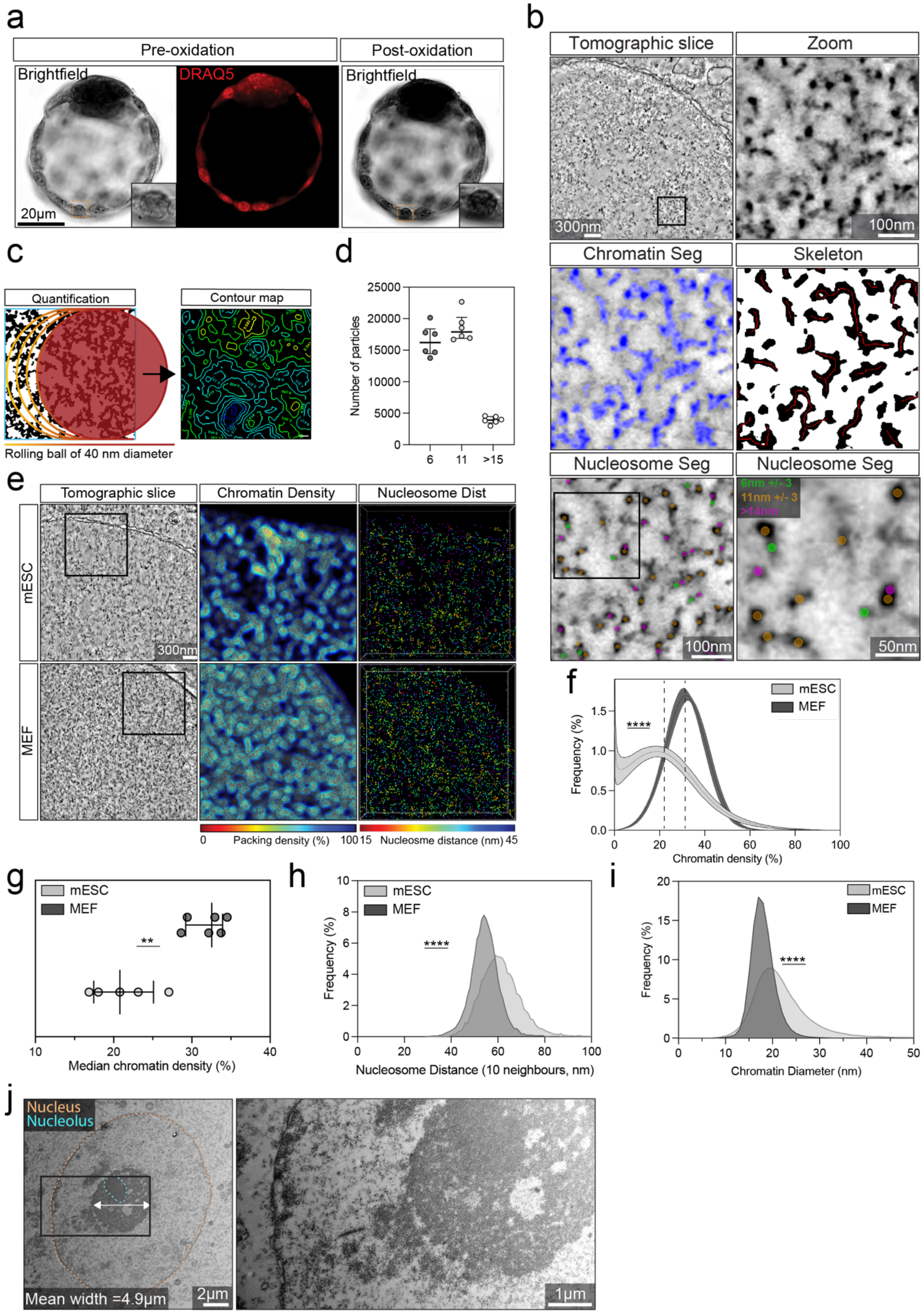

**Figure S1: Quantification pipeline for analysis reveals differences in chromatin ultrastructure.** a, Mouse blastocyst stained with DRAQ5 (red) before and after photo-oxidation. Inset shows zoom of a nucleus highlighting the polymerization of DAB, indicated by the presence of dark nuclei after oxidation observed by bright field. b, Segmentation of chromatin in a mouse 16-cell embryo. First panel shows a single tomographic slice, right shows zoom of the black box. Chromatin segmentation overlayed in blue, chromatin skeleton is shown in red over chromatin in black (middle panels). Bottom panel shows nucleosome segmentation with zoom on the right of area indicated by the black square. 5 nm densities are shown in green, 11 nm in pink, and >11 nm in orange. c, Cartoon with contour map of rolling ball-based quantification of chromatin packing. d, Quantification of the number of nucleosomes across the indicated size ranges. N=6 mouse embryonic fibroblast (MEF) cells across two biological replicates. e, Tomograms and quantitative maps of mouse embryonic stem cells (mESC) cultured in 2i LIF media and MEF, zoomed region (middle and right panels) indicated by the black box. Chromatin density, contour maps displaying chromatin packing density indicated by the color-coded scale bar. Nucleosome distance, nucleosomes colored based on their distance to 3 nearest neighbors indicated by the colored scale bar. f, Quantification of chromatin packing density, error bars show standard deviation. N= 5 mESCs, 6 MEF cells, 2 biological replicates. ****p <0.0001, Wilcoxon test. Dashed lines show the median value. g, Median chromatin packing density in the indicated cell types, each dot represents an individual cell used in f. Bars show median and interquartile range. **p <0.01, Mann-Whitney test. h, Quantification of the distance of nucleosomes to 10 nearest neighbors. N= 5 mESCs, 6 MEF cells, 2 biological replicates. ****p <0.0001, Wilcoxon test. i, Quantification of chromatin diameter in the indicated cell types. N= 5 mESCs, 6 MEF cells, 2 biological replicates. ****p <0.0001, Wilcoxon test. j, Transmission electron microscopy (TEM) images of a mouse 16-cell embryo, zoom to the right indicated by the black box, nucleus and nucleolus are highlighted by orange and blue dashed lines respectively. The mean width of nucleolar heterochromatin domains pooled from 12 nuclei is also shown.

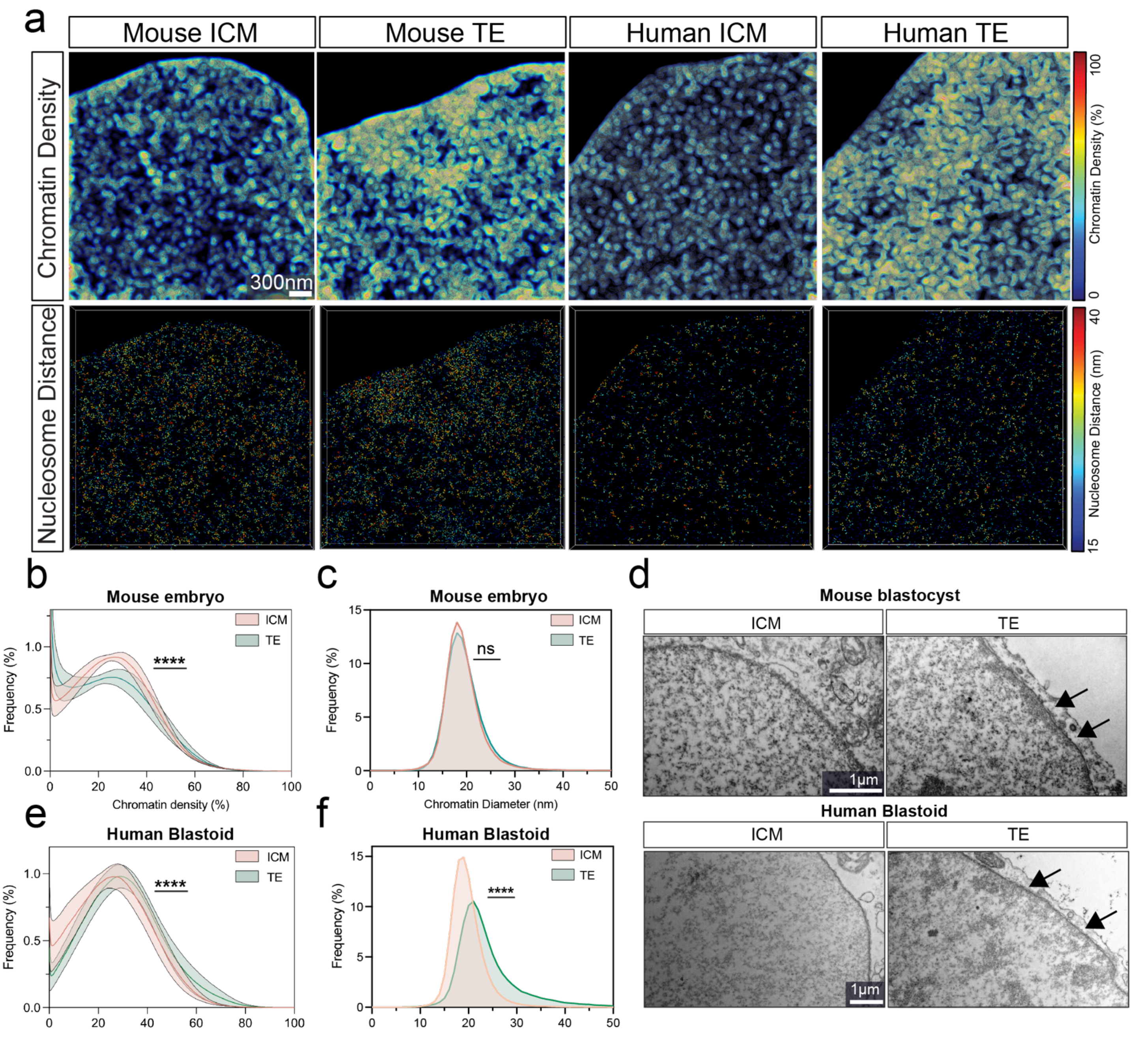


**Figure S2: Increased chromatin compaction is observed at the nuclear periphery of TE cells in mouse embryos and human blastoids.** a, Uncropped quantitative maps of the images shown in Fig. 2a. Chromatin density (top) and the distance of nucleosomes to three nearest neighbors (bottom) is indicated by the color-coded scale bars. b, Histogram of global chromatin density in ICM and TE cells of the mouse embryo, error bands show standard deviation. N = 7 nuclei from four embryos in two biological replicates. ****p <0.0001, Wilcoxon test. c, Quantification of chromatin diameter of the cells used in b. ns =non-significant, Wilcoxon test. d, Histogram of global chromatin density in ICM and TE cells of human blastoids, error bands show standard deviation. N = 9 nuclei from four blastoids in two biological replicates. ****p <0.0001, Wilcoxon test. e, Quantification of chromatin diameter of the cells used in d. ****p <0.0001, Wilcoxon test. f, Transmission electron microscopy (TEM) images of the indicated cell types. Arrows indicate heterochromatin at the nuclear membrane.

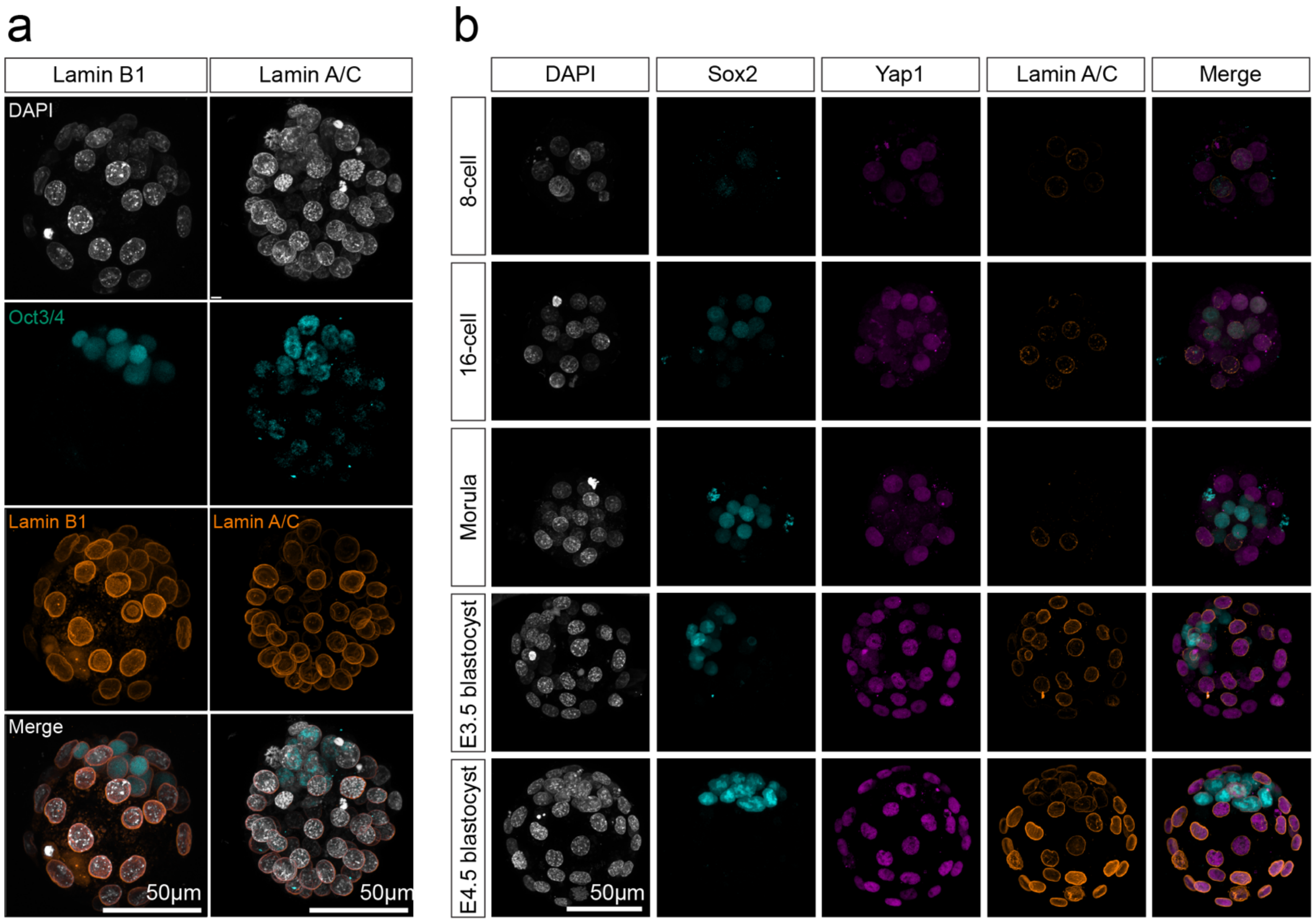


**Figure S3: Lamin A/C is upregulated in TE cells at the time of TE specification.** a, Immunofluorescence (IF) images of mouse blastocysts, with DAPI (grey), Oct3/4 labeling the ICM (cyan) and either Lamin A/C (orange), replicated data from Fig 3A or Lamin B1 (orange). b, IF images of mouse embryos at the indicated stages with DAPI (grey), Sox2 (cyan, ICM), Yap1 (magenta, TE), Lamin A/C (orange).

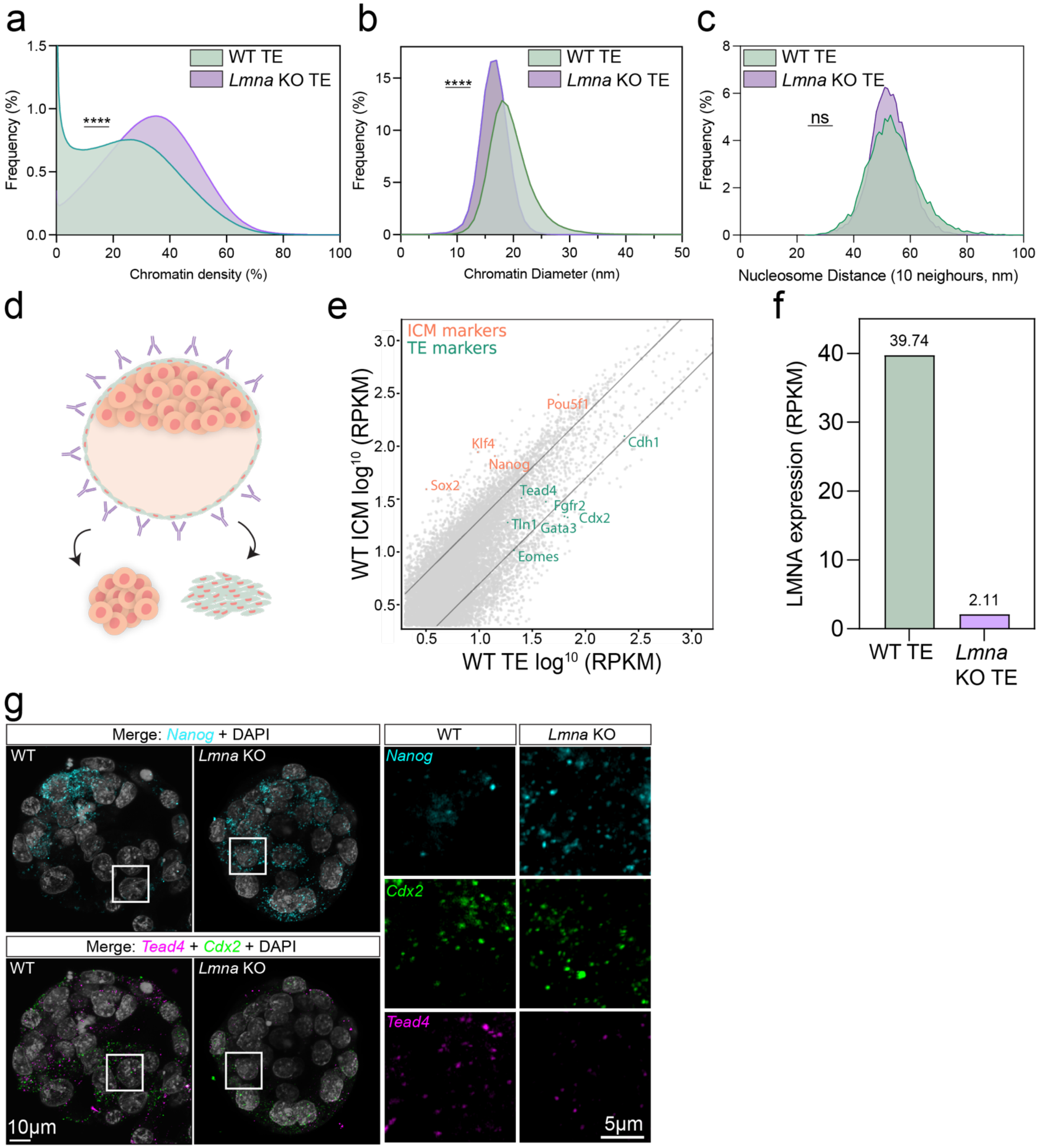


**Figure S4: Lamin A/C is specifically upregulated in TE cells at the time of specification.**

a, Quantification of global chromatin packing density in TE cells of WT and *Lmna* KO embryos. N= 7 cells across 4 embryos, two biological replicates. ****p <0.0001, Wilcoxon test. b, Quantification of chromatin diameter in TE cells of WT and *Lmna* KO embryos. N= 7 cells across 4 embryos, two biological replicates. ****p <0.0001, Wilcoxon test. c, Quantification of the average distance of nucleosomes to 10 nearest neighbors in TE cells of WT and *Lmna* KO embryos. N= 7 cells across 4 embryos, two biological replicates. ns=non-significant, Wilcoxon test. d, Cartoon of immunosurgery used to isolate ICM and TE cells from mouse blastocysts. e, Graph showing RNA sequencing counts in WT ICM and TE cells that were used in the RNA-seq experiments shown in Fig. 4j. Enriched expression for ICM genes (pink) in the ICM and TE genes (green) in the TE are highlighted. f, Quantification of *Lmna* RNA expression in WT and *Lmna* KO embryos from RNA-seq. g, RNA labeling of *Nanog* (cyan), *Cdx2* (green), and *Tead4* (magenta) in WT and *Lmna* KO blastocysts. Right panels show an overlay with DAPI (gray).

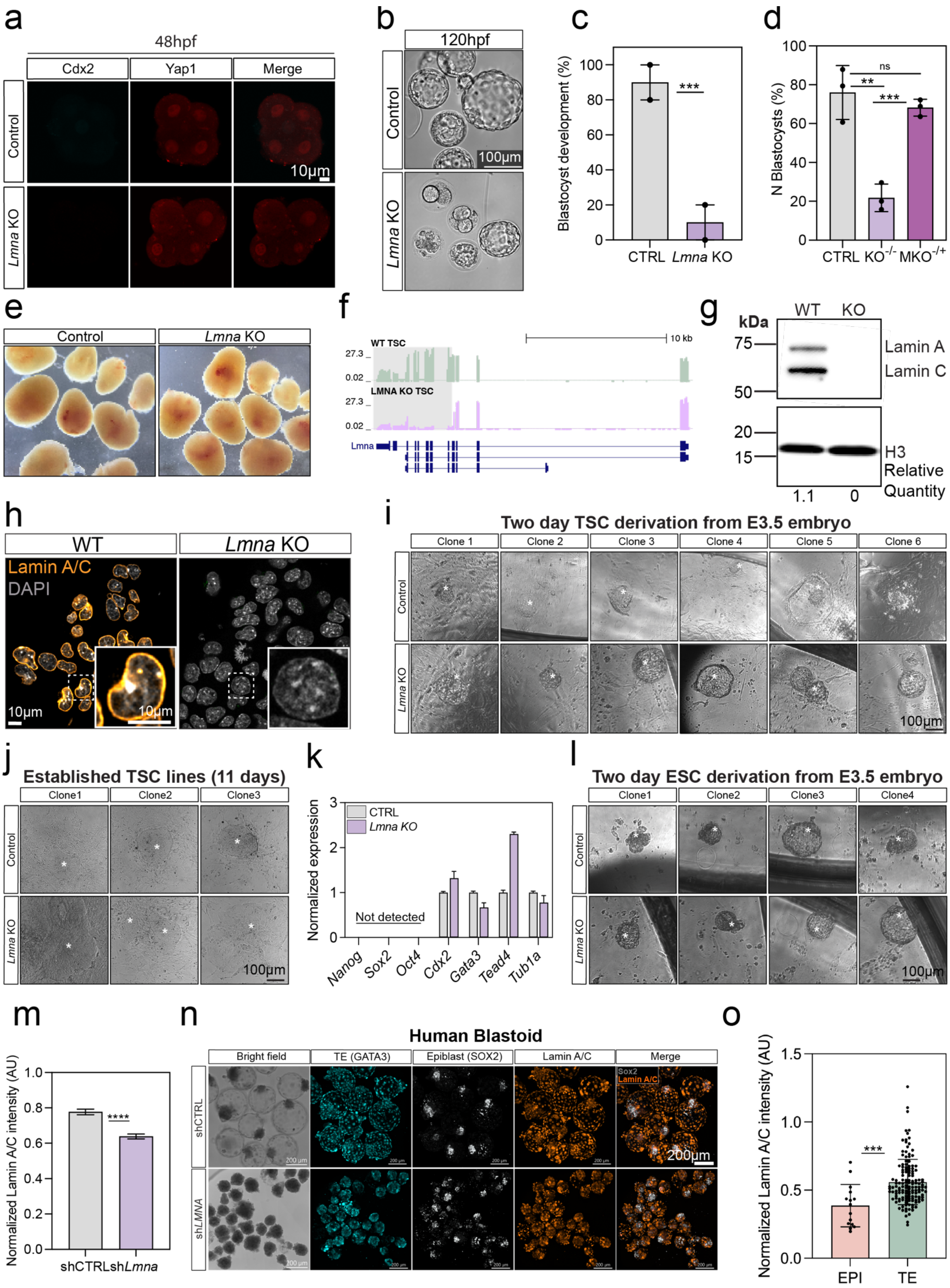
a
48hpf
Cdx2
Yap1
Merge
Control
Lmna KO
10µm
b
120hpf
Control
Lmna KO
100µm
c
Blastocyst development (%)
***
CTRL
Lmna KO
d
N Blastocysts (%)
ns
**
***
CTRL
KO-/-
MKO-/+
e
Control
Lmna KO
f
WT TSC
LMNA KO TSC
27.3
0.02
10 kb
Lmna
g
kDa
WT
KO
75
50
20
15
Lamin A
Lamin C
H3
Relative Quantity
1.1
0
h
WT
Lmna KO
Lamin A/C
DAPI
10µm
i
Two day TSC derivation from E3.5 embryo
Clone 1
Clone 2
Clone 3
Clone 4
Clone 5
Clone 6
Control
Lmna KO
100µm
j
Established TSC lines (11 days)
Clone1
Clone2
Clone3
Control
Lmna KO
100µm
k
Normalized expression
CTRL
Lmna KO
Not detected
Nanog
Sox2
Oct4
Cdx2
Gata3
Tead4
Tub1a
l
Two day ESC derivation from E3.5 embryo
Clone1
Clone2
Clone3
Clone4
Control
Lmna KO
100µm
m
Normalized Lamin A/C intensity (AU)
****
shCTRL
shLmna
n
Human Blastoid
Bright field
TE (GATA3)
Epiblast (SOX2)
Lamin A/C
Merge
Sox2
Lamin A/C
shCTRL
shLMNA
200µm
o
Normalized Lamin A/C intensity (AU)
***
EPI
TE

**Figure S5: Lamin A/C regulates blastocyst development in mouse embryos and human embryo models.**

a, Immunofluorescence (IF) images of control and *Lmna* KO embryos 48 hours post fertilization (hpf). Cdx2 (cyan) and Yap1 (red). b, Brightfield images of control and *Lmna* KO embryos 120 hpf. c, Quantification of the number of WT and *Lmna* KO blastocysts at 120 hpf. N=10 embryos, two biological replicates, ***p<0.001, Student's t-test. d, Rate of survival of control and *Lmna* KO embryos from Fig. 5d. Survival after chimeric *Lmna* KO$^{+/-}$ (depletion of *Lmna* KO in only one cell of a 2-cell embryo) is also shown. **p <0.01, ***p <0.001, ns= non-significant, one-way ANOVA. e, Images of decidua from control and *Lmna* KO embryos as quantified in Fig. 5f. f, RNA sequencing tracks of WT and *Lmna* KO trophoblast stem cells (TSCs) showing deletion of *Lmna* exon 1 and a second larger deletion between exons 4-10. g, Western blot of WT and *Lmna* KO TSCs, Lamin A/C and H3 (loading control) are shown. kDa= kilodaltons. Full uncropped Western blots using two different antibodies can be found in source data. h, IF images of WT and *Lmna* KO TSCs; Lamin A/C in orange, DAPI in gray. i-j, Images of WT and *Lmna* KO TSCs at the indicated times post induction. N= 6 WT and *Lmna* KO TSC clones. k, qPCR of control and *Lmna* KO TSCs. N= 1 biological replicate, 4 technical replicates.l, Images of WT and *Lmna* KO embryonic stem cells (ESCs) at the indicated stage post induction. N= 4 ICM ESC clones. m, Quantification of Lamin A/C intensity in shCTRL and sh*LMNA* KD blastoids. N=10 shCTRL blastoids (1014 TE cells) and N=31 sh*LMNA* blastoids (445 TE cells), ****p < 0.0001, Student's t-test, 2 biological replicates. n, IF images of shCTRL and sh*LMNA* human blastoids. Brightfield, Gata3 (cyan), Sox2 (gray), Lamin A/C (orange). o, Quantification of Lamin A/C intensity in epiblast (EPI) and TE cells of WT blastoids. N=10 blastoids. ***p < 0.001, Student's t-test, 2 biological replicates.

**Movie S1. 3D Segmentation of ChromEMT data** Chromatin is first segmented (blue) from a tomogram and then used to generate chromatin filaments (pink, see Materials and methods). Nucleosome-like densities are segmented in three size ranges 5nm (green), 10nm (orange), and >10nm (light pink). This video shows tracing of one chromatin region from the tomogram in Fig 1b.

**Movie S2. Segmentation of an individual chromatin chain** Chromatin (purple) and nucleosomes (orange) of a single chromatin chain. Other chromatin chains are shown in gray.
